# Non-Degenerate Two-Photon Imaging of Deep Rodent Cortex using Indocyanine Green in the water absorption window

**DOI:** 10.1101/2024.01.13.575485

**Authors:** Alankrit Tomar, Shaun A. Engelmann, Aaron L. Woods, Andrew K. Dunn

**Affiliations:** Department of Eletrical and Computer Engineering, The University of Texas at Austin, 2501 Speedway, Austin, TX 78712, USA; Department of Biomedical Engineering, The University of Texas at Austin, 107 W. Dean Keeton, Austin, TX 78712, USA

## Abstract

We present a novel approach for deep vascular imaging in rodent cortex at excitation wavelengths susceptible to water absorption using two-photon microscopy with photons of dissimilar wavelengths. We demonstrate that non-degenerate two-photon excitation (ND-2PE) enables imaging in the water absorption window from 1400-1550 nm using two synchronized excitation sources at 1300 nm and 1600 nm that straddle the absorption window. We explore the brightness spectra of indocyanine green (ICG) and assess its suitability for imaging in the water absorption window. Further, we demonstrate *in vivo* imaging of the rodent cortex vascular structure up to 1.2 mm using ND-2PE. Lastly, a comparative analysis of ND-2PE at 1435 nm and single-wavelength, two-photon imaging at 1300 nm and 1435 nm is presented. Our work extends the excitation range for fluorescent dyes to include water absorption regimes and underscores the feasibility of deep two-photon imaging at these wavelengths.

## 1. Introduction

Multiphoton fluorescence microscopy (MPM) is widely used to image cerebrovascular structure in the rodent brain with sub-micron resolution [1]. The applications of MPM in brain imaging include blood flow monitoring [2–4] imaging of neuronal activity [5], partial pressure of oxygen (pO_2_) characterization [6–8], assessing microvasculature post-vessel occlusion [9], to name a few. The technique relies on non-linear excitation of a fluorescent dye by two or more photons. Conventionally, the excitation photons are of the same wavelength and this process is referred to as degenerate two-photon excitation (D-2PE). The same excitation process can be induced by two photons of different wavelengths with two different excitation sources. This process is referred to as ND-2PE. We explore this concept to illustrate *in vivo* multiphoton imaging in the water absorption window.

ND-2PE has also been referred to as two-color excitation. In this work we adopt the abbreviation ND-2PE to denote this excitation process. Building on a theoretical framework proposed for non-degenerate excitation by McClain [10], one of the first experimental demonstration of ND-2PE involved two-photon excitation of an organic fluorophore by two different wavelengths [11]. Consequently, ND-2PE was used to demonstrate resolution enhancement [12] and a reduction in out-of-focus background [13]. Both outcomes were realized using two sources oriented at tilted angles relative to each other. Other ND-2PE studies demonstrated efficient optical sectioning [14], increased penetration depth [15] and even application in direct band-gap semiconductors [16]. None of these studies detailed usage of ND-2PE microscopy for biological samples. In 2009, the first ND-2PE based laser scanning microscope for acquiring UV fluorescence images of cells was developed [17]. Consequently, Mahou et al. demonstrated simultaneous imaging of three chromophores using ND-2PE in the mouse brain and spinal cord tissue labelled with fluorescent proteins [18]. Further work conducted on the same principle extended the research by illustrating large-volume ND-2PE imaging on fluorescent-labeled fixed brain tissue [19]. In 2016, Yang et al. established a theoretical foundation for comparing ND-2PE and D-2PE. Subsequently, they conducted empirical experiments to validate their theory, focusing on fluorescence signal levels, power dependence, and attenuation length within a fluorescein sample [20]. Building on their work, Sadegh et al. measured the non-degenerate excitation efficiency spectras relative to degenerate excitation for multiple dyes including fluorescien, SR101, EGFP and observed higher ratios than those predicted by theory [21, 22]. Strongari et al. and Ung et al. proposed using ND-2PE for Fluorescence Lifetime Microscopy (FLIM) of nicotinamide adenine dinucleotide(phosphate) (NAD(P)H) and flavin adenine dinucleotide (FAD) in reconstructed human skin and live *C*.*Elegans* [23, 24]. A computational framework involving numerical modeling of resolution and imaging depth for D-2PE and ND-2PE was carried out by Cheng et al. [25]. To our current understanding, the sole study showcasing *in vivo* rodent vascular imaging through ND-2PE has been carried out by Perillo et al [26]. In their research, they successfully exhibited that ND-2PE extends the penetration depth for *in vivo* vascular imaging by approximately 160 micrometers as compared with D-2PE. The work illustrated effective utilization of overlapping two excitation sources to expand the range of wavelengths available for imaging. Through our work, we aim to achieve the same objective but with a distinct focus on extending the excitation range to encompass the water absorption window.

The two- and three-photon excitation efficiency of a dye depends on its action cross-section at the excitation wavelength. In order to achieve high fluorophore brightness while still maintaining low excitation powers, the dye should ideally be excited at a wavelength at which its action cross-section is highest. Unfortunately, for certain dyes, the action cross-section coincides with wavelengths associated with high water absorption i.e. 1400-1550 nm. Imaging in this wavelength range is undesirable due to one photon absorption of excitation light by water molecules. As a result, the number of ballistic photons reaching the focal plane becomes negligible leading to low probability of non-linear excitation of fluorophores of interest and undesired tissue heating due to the water absorption. In this paper, we explore the feasibility of using ICG as a probe for *in vivo* 2-photon microscopy in the water absorption window. ICG is a fluorescent dye that is used clinically to visualize vasculature in surgery. The prevalence of ICG in human clinical surgeries and its low cost serve as a motivation to investigate its feasibility for *in vivo* ND-2P microscopy.

We show that two temporally and spatially synchronized, spectrally different, pulsed laser sources can be used to excite a dye at a wavelength that falls in the water absorption window if their wavelengths straddle the water absorption window. Prior to this, we investigate the relative excitation efficiency of ICG from a range of 1250-1600 nm. We then demonstrate an optical layout for ND-2PE using two excitation sources at 1300 nm and 1600 nm, respectively. These are used to excite ICG at an effective wavelength of 1435 nm, which lies within the water absorption window. We use the ND-2P imaging setup to image rodent cortex up to 1.2 mm in vivo. Finally, we also compare the performance of our ND-2P imaging with degenerate excitation at 1300 nm and the effective ND wavelength (1435 nm).

## 2. Materials and Methods

### 2.1. Ultrafast Excitation Sources

Two-photon imaging was performed using a custom-built fluorescence microscope with two separate excitation sources. Two noncollinear optical parametric amplifier lasers (NOPA, Spectra-Physics) were pumped by a single 1 MHz, 70 W pulsed laser (Spirit 1030-70, Spectra-Physics). Each NOPA was pumped at 35 W to ensure synchronization of the pulse trains. The two NOPA lasers were set to 1300 nm and 1600 nm, respectively. The measured spectra for the excitation sources are shown in Fig. 1. The red shaded region depicts the approximate water absorption window (detailed in section 3.1).

**Fig. 1.**
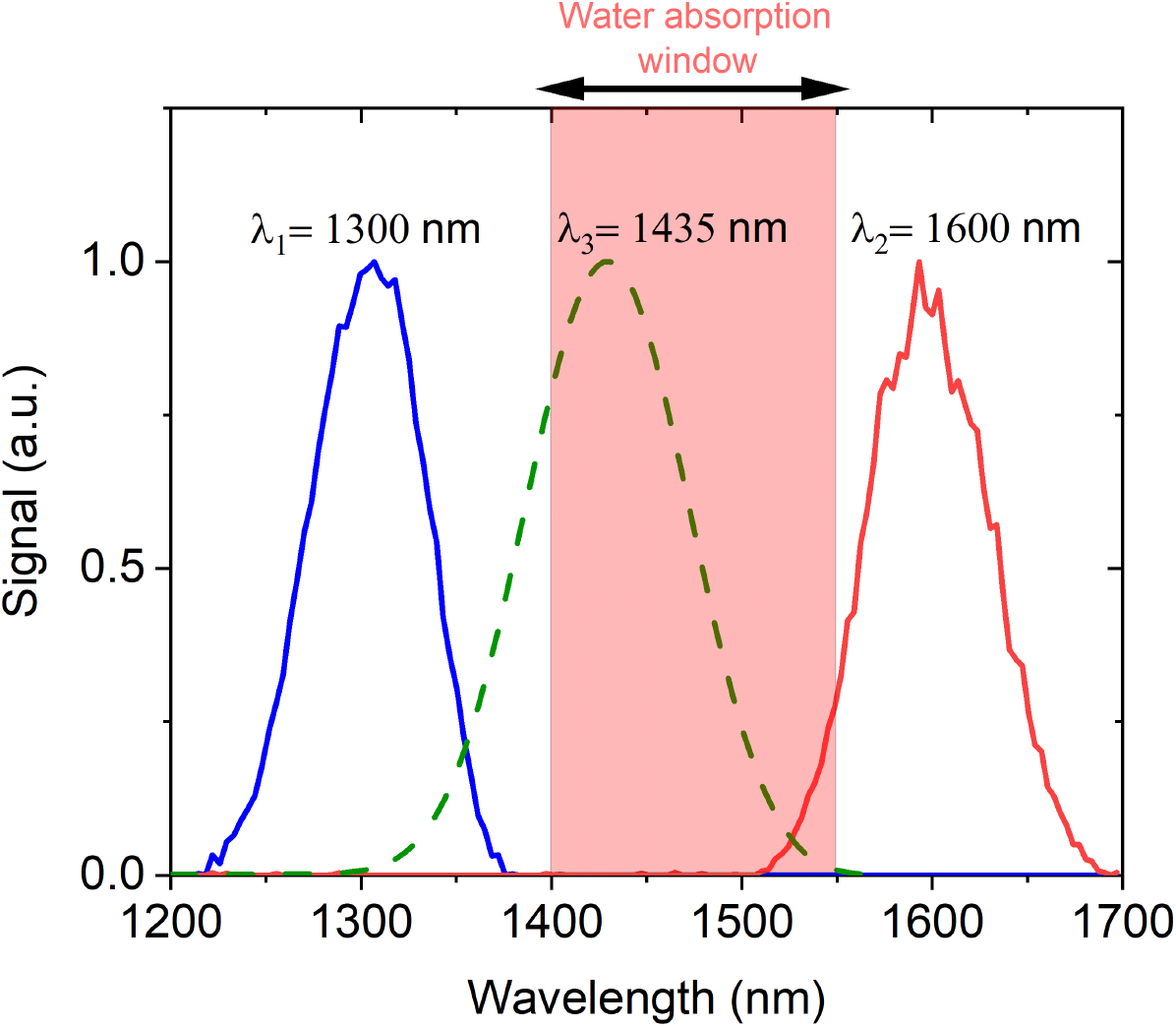
Measured spectra for the excitation sources (in blue and red), calculated spectra for ND effective wavelength (in green). The ND effective wavelength spectra was calculated from a cross-correlation between the measured spectra for the excitation sources

### 2.2. Multiphoton Microscope Setup

A home-built multiphoton microscope was used for imaging. Long pass filters (LPF1:FEL0850, Thorlabs, LPF2:FEL0850, Thorlabs) were used to remove any residual unwanted light from the NOPA laser outputs. A combination of half-wave plates (SAHWP05M-1700, Thorlabs for NOPA 1 and AQWP05M-980, Thorlabs for NOPA 2) and polarizing beamsplitters (PBS 124, Thorlabs) were used to control the power of excitation beams for imaging purposes. Long pass filters (LPF 3: FELH 1150, LPF 4: FF01-937) were used to restrict the excitation wavelengths to the desired values. Note that LPF 3 was added for the purpose of experiments discussed in section 3.2.1. This filter was not used for experiments discussed in section 3.2.2. Upon spatial and temporal alignment (discussed in section 2.2), the two excitation beams were directed to a pair of galvanometric scanners (QS7XY-AG, Thorlabs) for raster scanning. A scan lens (f=50 mm, SL50-3P, Thorlabs) and a Plössl tube lens were used in conjunction (f=200 mm, 2x AC508-400-C, Thorlabs) to expand the beam to fill the back aperture of a 25x objective (XLPLN25XSVMP2, 1.0 NA, Olympus). The objective focuses the excitation light onto the sample leading to emission of fluorescence signal. The emitted photons were reflected towards the detection path by a dichroic mirror (FF980-DI01-T1, Semrock) wherein they passed through a set of emission filters (893/209, Semrock, SP; 855/210, Semrock, SP) before getting collected by photo-multiplier tubes (H10770PB-50, Hamamatsu). The control of the stages, scanners, and image acquisition was carried out by a custom-made software in LabView. The image analysis was undertaken using Fiji [27]. A schematic of the microscope layout is shown in Fig. 2 (a).

**Fig. 2.**
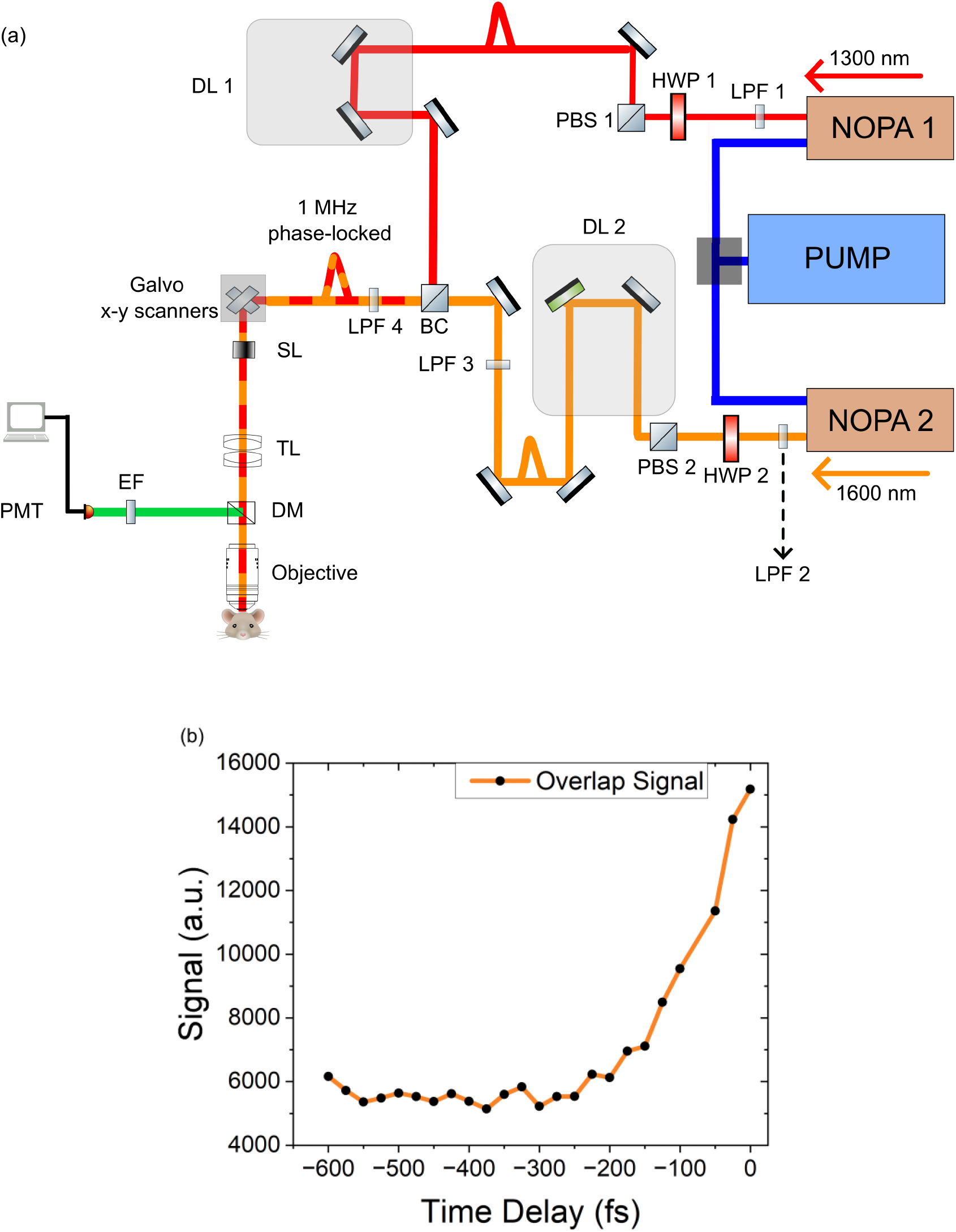
(a) Schematic of optical layout for aligning excitation sources spatially and temporally. NOPA 1 is overlapped in time relative to NOPA 2 using delay line 1 (DL1). (b) Non-degenerate excitation signal as a function of time delay between pulses of NOPA 1 and NOPA 2. NOPA: noncollinear parametric amplifier, LPF: long pass filter, HWP: half wave plate, PBS: polarizing beam splitter, DL: delay line, C: collimators BC: beam combiner, SL: scan lens, TL: Tube lens, DM: Dichroic Mirror, EF: emission filters, PMT: photomultiplier tube

### 2.3. Optical Layout for Temporal and Spatial Alignment

For the non-degenerate excitation process to occur efficiently, it is necessary that pulses from one excitation source align spatially and temporally at the focal plane with the pulses from the second excitation source. The optical layout shown in Fig. 2(a) was designed to spatially and temporally align the pulses from the two excitation sources, NOPA 1 and NOPA 2, at the imaging plane of the microscope.

The spatial alignment is performed with the help of steering mirrors, collimators and pinholes in the paths of the two beams. First, the beams are roughly aligned at the beam combiner (BC). Consequently, two pinholes, 1.5 m apart and placed after BC, are used to laterally align the beams precisely over one another. Collimators in the path of the beams were adjusted to ensure relative axial alignment. To verify coarse spatial alignment in the focal plane, we imaged a lens cleaning paper soaked with ICG, our dye of interest. Adjustments were made to one of the mirrors in the path of NOPA 1 to make the images coincident.

For the beams to align temporally, it was necessary that pulses travel the same path length from the pump laser to the imaging plane. As a first step, the manual stage (DL 2) in the path of NOPA 2 is moved as to roughly get the path lengths equal. For precise alignment, we set up a cuvette well with 100 *μ*L of ICG solution and measure the fluorescence signal originating from the sample as a function of the position of a piezo stage (DL 1) in the path of NOPA 2. As the stage moves through its range, it adjusts temporal delay in the path of NOPA 2 and at a specific position of the stage, when the pulses are precisely aligned, we measure a peak in the signal from the dye sample. The peak signifies the non-degenerate signal arising as a result of spatial and temporal overlap of the pulses in the focal plane. Figure 2(b) depicts the fluorescence signal as a function of temporal delay in the path of NOPA 2. As a result of the overlap of the two pulses at wavelengths *λ*_1_ and *λ*_2_, we get effective excitation at a wavelength of *λ*_3_ = (2*λ*_1_*λ*_2_)/(*λ*_1_ + *λ*_2_). In our setup, by using excitation sources *λ*_1_=1300 nm and *λ*_2_=1600 nm, the effective ND excitation is achieved at *λ*_3_=1435 nm. The spectra of the effective wavelength is shown in green in Fig. 1 and is measured via cross-correlation between the two excitation sources. The peak signal obtained in Fig.2(b) is optimized with slight adjustments to the spatial alignment. The position of the piezo stage is noted and is maintained through the experimental procedure. The spatial and temporal alignment process was verified before every experiment.

### 2.4. Characterization of Indocyanine Green

To identify wavelengths that would be feasible for 2-photon imaging of ICG, we analyzed its brightness spectra from 1250 nm to 1600 nm. We dissolved ICG (Sigma Aldrich) in a 5% bovine serum albumin-phosphate buffered saline solution to prepare a 100 *μ*M dye solution. A cuvette well was filled with 0.75 ml of the 100 *μ*M solution and a cover slip was slid carefully onto the well avoiding any air bubbles in the sample. Heavy water was used as immersion media for the objective. The average power was measured and was maintained at 6 mW at the focal plane for all wavelengths during the experiment. To avoid any absorption of excitation photons due to water content in the sample, we found the top of the sample and imaged the same field of view for all wavelengths. A segment located at the same position within a 3-frame averaged image was selected for each wavelength. Following which the mean intensity of the segment was attributed as signal for that specific wavelength.

### 2.5. Animal Preparation and Imaging

Animal procedures were approved by The University of Texas at Austin Institutional Animal Care and Use Committee. Mice were surgically fit with a 5 mm cranial window under the influence of a anaesthetic agent. The temperature of the mouse was maintained at 37.6 °C during the craniotomy procedure. The mice recovered for at least 2 weeks before imaging. Powdered ICG was dissolved in 5% bovine serum albumin-phosphate-buffered saline (PBS) solution to a concentration of 1 mg/ml. The blood plasma of the vascular network in the cortex was labelled through a 100 *μ*L retro-orbital injection of the ICG solution. During imaging, the average power incident on the surface was increased with depths but did not exceed 91 mW.

## 3. Experimental Results

### 3.1. Brightness curve for Indocyanine Green

We mapped the brightness spectra of ICG from 1250 to 1600 nm at intervals of 50 nm, as shown in blue in Fig. 3. The spectral shape of the brightness spectra have been demonstrated to match reported action cross-section of dyes [28]. The brightness curve was normalized with respect to the peak brightness at 1450 nm.

**Fig. 3.**
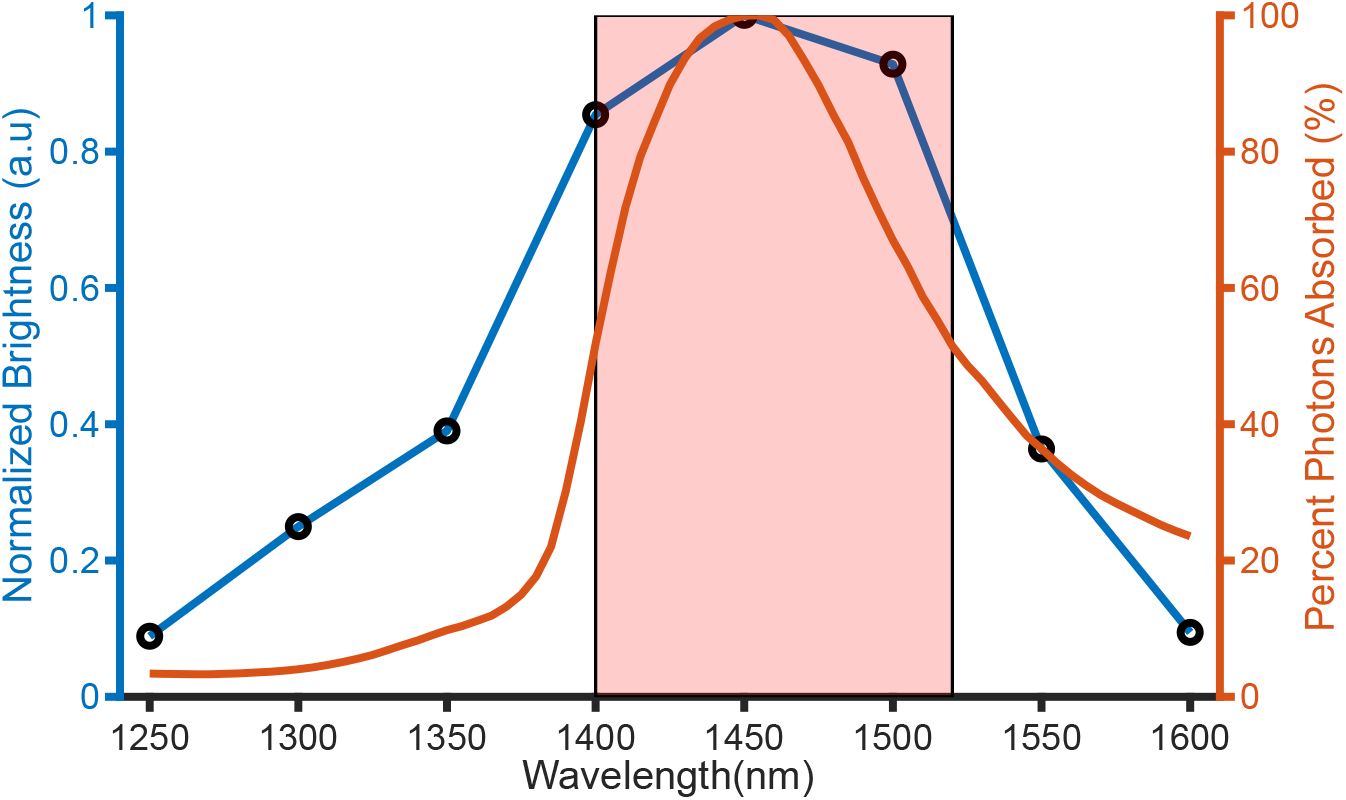
*Left*: Two-Photon Brightness Spectra for ICG, *Right*: Percent of excitation photons absorbed by water at a depth of 1 mm in brain tissue; The red box covers wavelengths at which 50% of the photons are absorbed by water at a depth of 1 mm

The red curve in Fig.3 illustrates the effect of absorption of ballistic photons by water in tissue. The absorption coefficients for water [29] were scaled by the water content in brain assumed to be 75% [30], and attenuation of the excitation light due to one photon absorption was assumed to follow simple Beer’s Law attenuation [31]. Further, the shaded area in red indicates wavelengths wherein more than 50% of the photons are absorbed by the tissue over a 1 mm pathlength. These wavelengths, ∼1400-1550 nm, are not suitable for imaging as excessive water absorption leads to tissue damage, which is undesirable. Note that the photons absorbed by water at 1300 nm and 1600 nm are approximately 95% and 80% less, respectively, than the photons absorbed at 1450 nm. This serves as a motivation for using 1300 nm and 1600 nm as excitation sources to excite ICG through ND-2PE at an effective wavelength of 1435 nm.

### 3.2. Non-degenerate two-photon in vivo imaging

*In vivo* imaging of the vascular structure of the mouse cortex was carried out using the two excitation sources at 1300 nm and 1600 nm. Due to rapid clearance of ICG, we could only investigate a few distinct slices of vascular structure rather than a continuous stack. At every depth investigated, 5 frames were averaged to produce a slice of 410 *μ*m x 410 *μ*m field of view (FOV). Sections 3.2.1 and 3.2.2 detail the *in vivo* experiments carried out using ND-2P using 1300 nm and 1600 nm excitation light. The vascular images shown in Fig. 4 and Fig. 5 are displayed on the same scale of pixel intensity values. The mean SNR in Fig. 4 was calculated using four sample vessels at each depth. Similarly, 2 vessels at the surface and 3 vessels at all other depths were used for mean SNR calculations in Fig. 5.

**Fig. 4.**
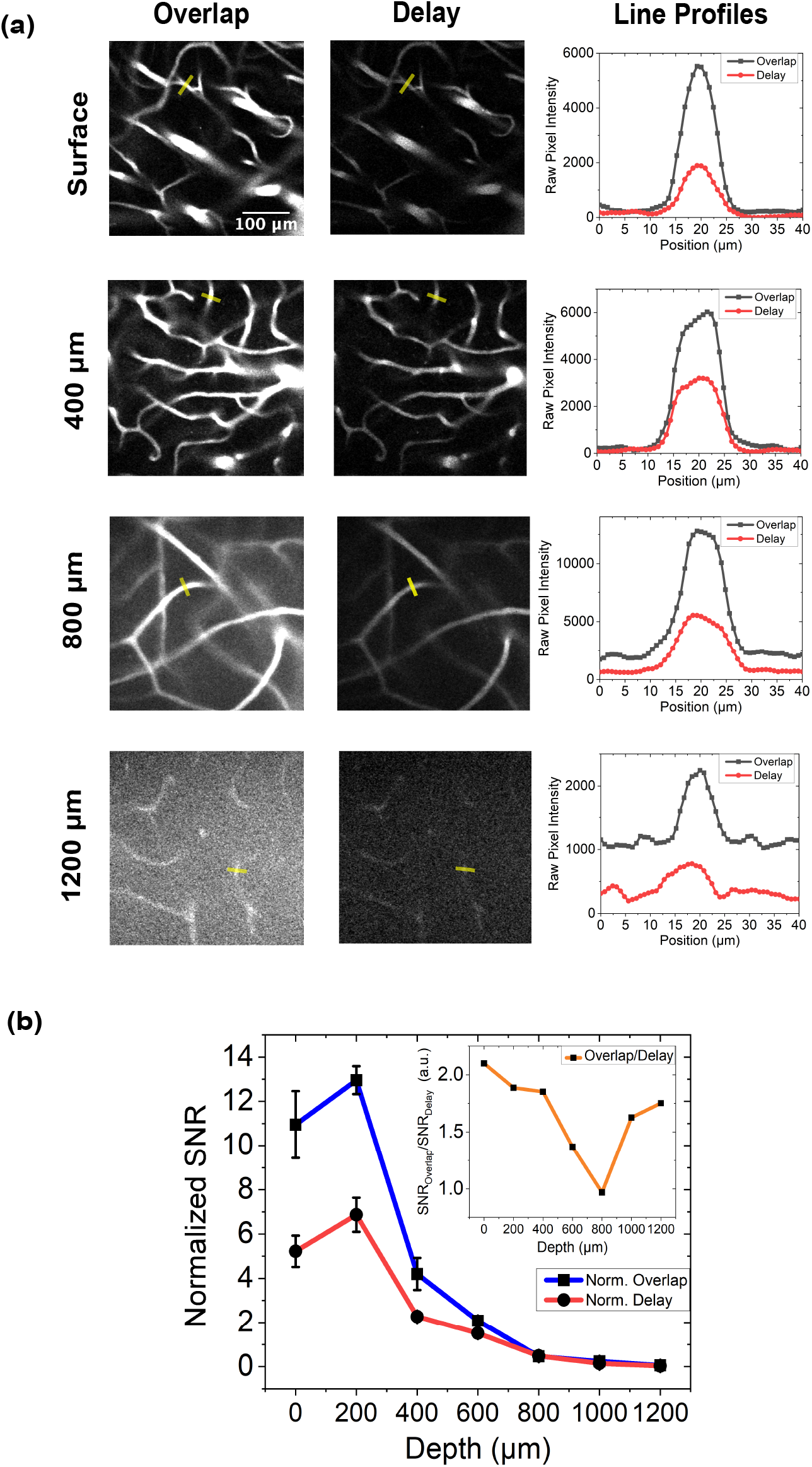
Deep *in vivo* two-photon images of ICG in rodent cortex blood vessels with ND-2PE. (a) 5-frame averaged x-y slices and raw intensity profiles of vessels at different depths under *Overlap* and *Delay* excitation schemes. (b) Mean SNR with standard error normalized to average power as a function of depth, *Inset*: Ratio of mean SNRs for *Overlap* and *Delay* conditions as a function of depth

**Fig. 5.**
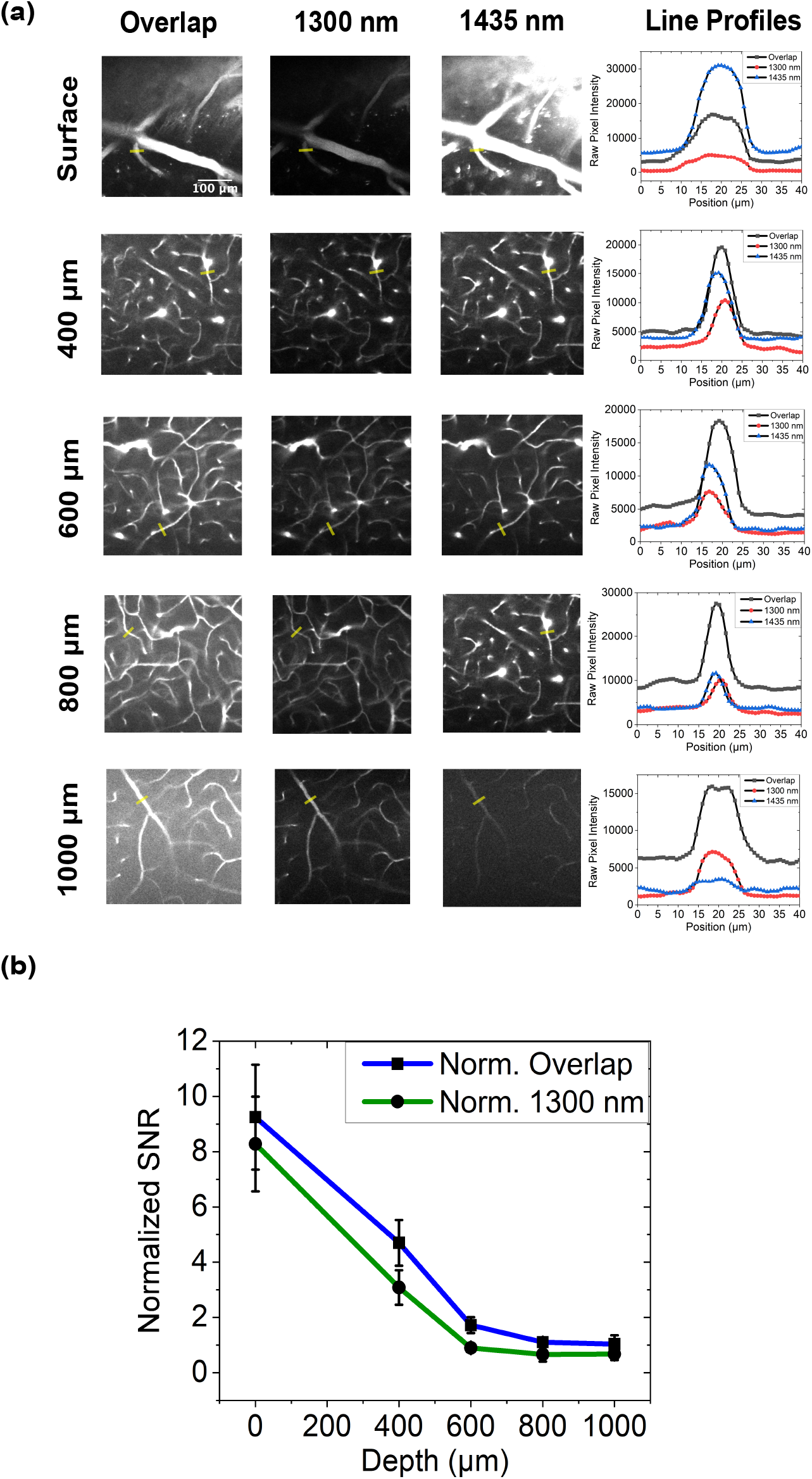
Two-photon images of blood vessles obtained with ND-2PE and degenerate excitation at 1300 nm and 1435 nm. (a) 5-frame averaged x-y slices and raw intensity profiles of vessels at different depths under *Overlap* and degenerate conditions (b) Mean SNR with error bars normalized to average power as a function of depth for *Overlap* condition and degenerate excitation at 1300 nm

#### 3.2.1. Comparison of *Overlap* and *Delay* excitation schemes

The *in vivo* imaging procedure was carried out with two different excitation schemes: (i) in which the two excitation pulses were spatially and temporally aligned at the focal plane; (ii) in which the pulses were delayed by 1000 fs. Henceforth, in the paper, we will refer to the two schemes as *Overlap* and *Delay*, respectively. We imaged the rodent cortex from a depth of 1200 *μ*m with slices taken at every 200 *μ*m up to the surface. The maximum average power incident on the brain surface was restricted below 91 mW. Note that the average power incident for the *Overlap* and *Delay* excitation schemes was the same at each depth.

Figure 4(a) illustrates the 2-D slices acquired at different depths with the two excitation schemes. Line profiles for certain vessels, marked by yellow bars, are also shown in Fig. 4(a). Although we acquired images at intervals of 200*μ*m, we display images in Fig.4(a) only at intervals of 400 *μ*m to make the figure concise. Figure S1 (a) depicts the slices at the remaining depths. Figure 4(b) depicts the normalized signal-to-noise (SNR) ratio for the different depths, respectively, with the two schemes. The SNR at each depth was normalized with respect to the average power used at that depth. The SNR is quantified as (*μ*_sig_-*μ*_bg_)/σ_bg_ where *μ*_sig_ denotes the mean signal intensity, *μ*_bg_ and σ_bg_ denote the mean and standard deviation of the background intensity, respectively. Table 1 depicts the mean SNR along with standard error for the *in vivo* experiments in Fig. 4. Figure S1(b) shows the normalized SNR curves on a logarithmic scale.

**Table 1.** Comparison of mean SNR (with standard error) normalized to average power at each depth for *Overlap* and *Delay* conditions for different depths.

| Depth ($\mu$) | Overlap | Delay |
| --- | --- | --- |
| Surface | 10.95 ± 1.51 | 5.22 ± 0.70 |
| 200 | 12.96 ± 0.63 | 6.87 ± 0.77 |
| 400 | 4.2 ± 0.73 | 2.27 ± 0.15 |
| 600 | 2.08 ± 0.16 | 1.52 ± 0.04 |
| 800 | 0.47 ± 0.06 | 0.48 ± 0.01 |
| 1000 | 0.24 ± 0.03 | 0.15 ± 0.00 |
| 1200 | 0.1 ± 0.01 | 0.03 ± 0.00 |

#### 3.2.2. Comparison of *Overlap* with degenerate excitation at 1300 nm and 1435 nm

While the previous results demonstrated the comparison between *Overlap* and *Delay* schemes, it was important to investigate if imaging under the *Overlap* condition was better than imaging with degenerate excitation. We evaluated how the ND excitation fared in comparison to imaging with degenerate excitation at 1300 nm and at 1435 nm, the effective ND wavelength, given by *λ*_3_ = (2*λ*_1_*λ*_2_)/(*λ*_1_ + *λ*_2_), where *λ*_1_=1300 nm and *λ*_2_=1600 nm.

Figure 5 (a) depicts the vascular images, along with line profiles, obtained with *Overlap*, degenerate excitation at 1300 nm, and degenerate excitation at 1435 nm. Further, Fig. 5(b) shows the average power-normalized SNR for the different depths. In order to facilitate a fair comparison between the three cases, we ensured that the average power at a specific depth remained consistent across all three cases.The average power incident at the highest depth was 46 mW. Blood vessels starting from 1000 *μ*m up to the surface with intervals of 200 *μ*m were investigated. We noticed significant damage to vascular structure at shallow depths of around 200 *μ*m. We discuss the possible cause for damage in the Discussion section. The images obtained at 200 *μ*m are shown in Fig. S2(a) in the supplementary materials. Further, Fig. S2(b-c) displays the normalized SNR curves on a linear and logarithmic scale, respectively.

## 4. Discussion

One of the first steps towards undertaking MPM entails choosing a suitable dye for imaging, and then an choosing an optimal excitation wavelength for that specific dye. The decision regarding dye selection, in most cases, depends upon the structure of interest. For example, vascular structures can be imaged by labeling the blood plasma [32] whereas functional imaging can be performed using calcium indicators like GCaMP [33]. On the other hand, choosing an excitation wavelength presents with a trade-off between the excitation efficiency of the dye and the brain tissue scattering and absorption properties [31]. For fluorescent dyes, the two photon absorption cross section dictates how well the dye absorbs excitation light as a function of wavelength. Further, water content in brain tissue is responsible for one photon absorption of the majority of ballistic photons at long excitation wavelengths. Ideally, an optimal excitation wavelength is at which the two photon absorption cross section for a dye is maximum and the water absorption and tissue scattering are minimum. From a practical perspective, a balance between 2P absorption cross section and the water absorption needs to be made for choosing an excitation wavelength. Thus, water absorption restricts the range of excitation wavelengths that are feasible for *in vivo* multiphoton imaging.

Our goal with this work was to demonstrate that the wavelengths susceptible to water absorption, namely 1400-1550 nm, could still be utilized for MPM using ND-2PE. In order to test this, we sought to investigate dyes that have high action cross-sections in the water absorption window. Although there have been detailed analyses of excitation efficiency of dyes from 1200-1600 nm [28], most of them do not investigate the dyes in the water absorption window. In 2017, Miller et al. demonstrated relative action cross-sections for ICG from 1200-1450 nm with a peak in efficiency at 1450 nm [34]. Further, Wang et al. showed that even though the absolute action cross-sections for ICG decreased from 1600 nm to 1800 nm [35], in vivo D-2P imaging at 1700 nm showed promise [36]. These studies motivated us to look into a detailed excitation efficiency spectra of ICG from 1250-1600 nm, which includes the water absorption window. Towards that goal, we investigated brightness spectra for ICG from 1250 nm to 1600 nm. Kara et al. note that brightness curves could vary from between imaging systems but provide sufficient information for selecting an optimal excitation wavelength [28]. It was found that the peak of the brightness curve for ICG coincides with the water absorption peak. Under traditional two-photon excitation schemes, imaging ICG at the peak of its brightness curve would be unfeasible due to high water absorption.

We propose that the two-photon excitation of ICG at its peak brightness wavelength could be initiated with the help of two photons of different wavelengths, namely ND-2P. As noted previously, under this excitation scheme, two photons of different wavelengths, *λ*_1_=1300 nm and *λ*_2_=1600 nm, can excite ICG at a third effective wavelength, *λ*_3_= 1435 nm, where *λ*_3_ is given by *λ*_3_ = (2*λ*_1_*λ*_2_)/(*λ*_1_ + *λ*_2_). The formula illustrates that two photons of 1300 nm and 1600 nm provide the same amount as energy as two photons of 1435 nm each. Moreover, the motivation for selecting those specific excitation wavelengths arises from the presence of local minima in the water absorption curve around these wavelengths [31]. Therefore, by straddling the water absorption window, these two wavelenghts can excite ICG more efficiently while avoiding most of the water absorption.

For the ND excitation process to be efficient, the point spread functions (PSFs) of the excitation sources need to be spatially and temporally aligned in the focal plane. Figure 2 details the optical setup used for the non-degenerate alignment procedure. The occurrence of the non-degenerate signal at 1435 nm is reliant on the precise overlap of the excitation beams in the focal plane. All *in vivo* experiments were undertaken only once the alignment was confirmed.

Once the brightness curves confirmed a peak for ICG in the water absorption range, we wanted to demonstrate ND excitation *in vivo*. Figure 4 illustrates results from *in vivo* imaging of rodent vascular structure from the surface of the cortex to about 1.2 mm. We sought to compare the obtained images between when the excitation pulses were temporally overlapped, *Overlap* condition, and when they are delayed with respect to one another, *Delay* condition. The pulses from the two excitation sources were delayed with respect to each other by ∼1000 fs. Line profiles in Fig. 4(a) show that the absolute signals from specific vessels were greater for the *Overlap* condition across all depths. It is pertinent to note that when the pulses are delayed, the detected signal is the sum of degenerate signal from 1300 nm and 1600 nm excitation light. On the other hand, when the pulses are overlapped, in addition to the degenerate excitation signal from the two sources, there is signal as a result of ND-2PE at an effective wavelength of 1435 nm. As this wavelength corresponds to a peak in the brightness curve for ICG, we get a high signal enhancement.

To quantify the differences further, we investigated SNRs, normalized by average power at all depths, for the two conditions. As shown in Fig. 4(b), the normalized SNR for both the conditions reduces with depth. The downward trend in both cases can be attributed to the fact that as depth increases, the out-of-focus background increases leading to a lower SNR. At shallower depths, the SNR is significantly higher for the *Overlap* condition compared to the *Delay* condition. This is due to considerably high absolute signal values in the *Overlap* than in the *Delay* condition. However, as the depth increases, the difference becomes less pronounced. The inset in Fig. 4 highlights that even at highest depths, the mean SNR for the *Overlap* condition is more than 1.5 times the SNR for the *Delay* condition, the exception being at 800 *μ*m wherein the SNRs become almost equal. Figure S2, which displays the SNR on a logarithmic scale, also illustrates higher SNR for *Overlap* than *Delay* at deeper depths. A higher SNR with ND excitation practically enabled access to depths beyond 1.2 mm but imaging further posed a risk of the objective hitting the cranial window. Hence, we restricted our highest imaging depth to 1.2 mm. After obtaining the 2D slices, we reinspected each depth to look for signs of tissue damage. Finding no evidence of damage, we concluded that non-degenerate imaging does not harm the vascular structure, despite exciting a fluorescent dye at a wavelength within the water absorption window. The results discussed above strongly support the validity of the proposed technique.

The brightness curve for ICG indicates that other than the optimal wavelength of excitation at 1450 nm, there are other candidate wavelengths for imaging. For example, as depicted in Fig. 3, despite the ICG fluorescence signal at 1300 nm being 75 percent less bright than at 1450 nm, photons at 1300 nm could penetrate deeper into the tissue due to lower water absorption [34]. Thus, we investigated how degenerate 1300 nm excitation would compare with the *Overlap* condition. Sadegh et al. have experimentally noted that the non-degenerate two photon action cross section (ND-TPACS) i.e. the action cross section at a particular wavelength associated with excitation via a non-degenerate process, is greater than the TPACS at the same wavelength with excitation via a degenerate process [21]. This is known as resonance enhancement effect. The effect implies that excitation at 1435 nm via ND-2PE with 1300 nm and 1600 nm should be more efficient than degenerate excitation 1435 nm. Although degenerate imaging at 1435 nm can be harmful to the tissue, there was motivation in verifying the results by Sadegh et al. *in vivo*.

Figure 5 shows the images obtained at different depths under the three excitation schemes: ND excitation with 1300 nm and 1600 nm, degenerate excitation at 1300 nm, and degenerate excitation at 1435 nm. Maintaining the same average power at the surface for a specific depth ensures a fair comparisons between the schemes. It was noted from line profiles in Fig. 5(a) that the absolute signal values for D-2PE at 1435 nm was higher compared to the other two excitation schemes at the surface. Subsequently, as depth increases, the signal values for D-2PE at 1435 nm reduces significantly. This is expected as 1435 nm ballistic photons suffer from high water absorption as depth increases leading to lower photons available at the focus to initiate the two-photon excitation process. It is worth noting that if the results by Sadegh et al. held true then the absolute signal values at the surface with the *Overlap* condition should have been more than that with D-2PE at 1435 nm. We have no reasons to believe that the results by Sadegh et al. do not hold true but there are a couple of possibilities why we observe a lower absolute signal with ND-2PE: (i) Sadegh et al. validated their results in cuvette samples and there is a possibility that the results do not hold as well for biological samples; (ii) The alignment of 1300 nm and 1600 nm in our system might not be as precise as that required for the resonance enhancement theory to hold true. Note that this discrepancy has little practical implications for *in vivo* excitation of ICG as imaging with D-2P at 1435 nm is unfeasible due to high water absorption.

For the reasons discussed above, we only compare the SNR values for the *Overlap* condition and D-2PE at 1300 nm. Figure 5(b) depicts that the SNR, normalized to average power used at each depth, for the *Overlap* condition, was found to be greater than that with degenerate excitation at 1300 nm for all depths. The downward trend in both graphs can be attributed to increased out-of-focus background at higher depths, which contributed to higher noise levels. Table 2 contains normalized SNR values which are depicted in Fig. 5(b). Figure S2 illustrates that the normalized SNR for degenerate excitation at 1435 nm outperforms the *Overlap* condition at 400 *μ*m. The consequences of imaging via D-2PE in the water absorption window became evident at shallower depths of 200 *μ*m, where visible damage to the vascular network was observed. The damage was likely caused due to absorption of ballistic photons by water while imaging deeper depths in the cortex. Figure S2(a) shows the line profiles and x-y slices at 200 *μ*m.

**Table 2.** Comparison of mean SNR (with standard error) normalized to average power at each depth for *Overlap* condition and degenerate excitation for 1300 nm at different depths.

| Depth ($\mu$) | Overlap | 1300 nm |
| --- | --- | --- |
| Surface | $9.25 \pm 1.9$ | $8.28 \pm 1.72$ |
| 400 | $4.70 \pm 0.83$ | $3.08 \pm 0.63$ |
| 600 | $1.72 \pm 0.29$ | $0.9 \pm 0.07$ |
| 800 | $1.10 \pm 0.14$ | $0.66 \pm 0.26$ |
| 1000 | $1.04 \pm 0.32$ | $0.68 \pm 0.22$ |

As noted previously by Miller et al., and confirmed in this study, ICG levels in the vascular network significantly decrease within 30 minutes from injection [31]. Thus, the *in vivo* images obtained in this work were restricted to 2D slices at specific depths, which serves as a limitation of our study. In future work, we hope to address this limitation by making use of specific variants of ICG [37] which have higher clearance time, enabling 3D vascular analysis. A three-dimensional structure of the brain is important to understand the structure of the brain in healthy [38, 39] and diseased mice [40].

## 5. Conclusion

MPM is limited in its ability to image fluorescent dyes exhibiting high excitation efficiency in the water absorption windows. We map the brightness spectra for ICG for a large range of wavelengths, including the water absorption window. We demonstrate that ND-2PE via two wavelengths beyond the water absorption window, can effectively excite ICG within the absorption window. The technique is capable of damage-free, *in vivo* imaging of vascular structures as deep as 1.2 mm in the rodent cortex. In conclusion, our research demonstrates the unique concept of circumventing tissue heating due to water absorption by utilizing non-degenerate excitation. This breakthrough opens new possibilities for using dyes for *in vivo* at wavelengths prone to water absorption.

## Data Availability Statemen

Data underlying the results presented in this paper are not publicly available at this time but may be obtained from the authors upon reasonable request.

