## Supplementary material for "Non-Degenerate Two-Photon Imaging of Deep Rodent Cortex using Indocyanine Green in the water absorption window" (from another version of this preprint: version 1, 2024-01-15)

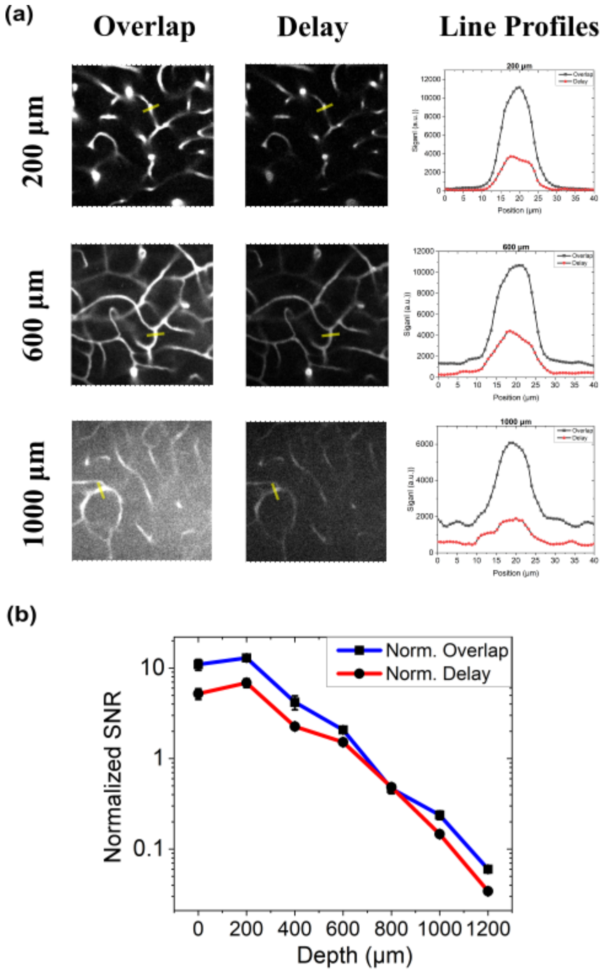


Figure S1: *In vivo* images using *Overlap* and *Delay* condition. (a) 5-frame averaged x-y slices and line profiles. (b) Normalized SNR on a logarithmic scale, as a function of depth

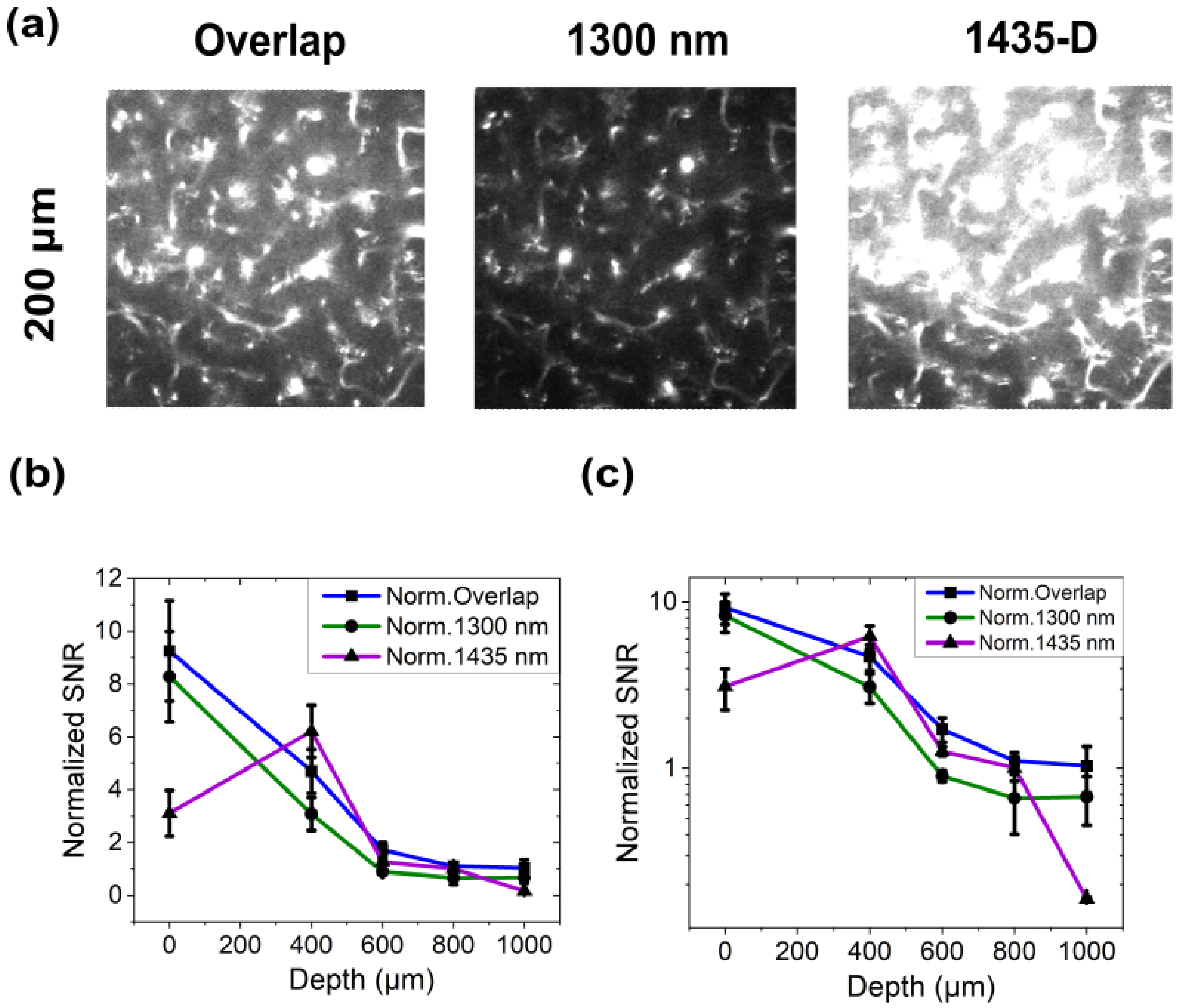


Figure S2: (a): *In vivo* images showing damage at 200 µm (a) 5-frame averaged x-y slices. (b) Normalized SNR on a linear scale for Overlap, D-2PE at 1300 nm and 1435 nm, as a function of depth. © (b) Normalized SNR on a logarithmic scale for Overlap, D-2PE at 1300 nm and 1435 nm, as a function of depth
